# Dual RNA-Seq analysis of SARS-CoV-2 correlates specific human transcriptional response pathways directly to viral expression

**DOI:** 10.1101/2021.02.09.430517

**Authors:** Nathan D Maulding, Spencer Seiler, Alex Pearson, Nick Kreusser, Josh Stuart

## Abstract

The SARS-CoV-2 pandemic has challenged humankind’s ability to quickly determine the cascade of health effects caused by a novel infection. Even with the unprecedented speed at which vaccines were developed and introduced into society, identifying therapeutic interventions and drug targets for patients infected with the virus remains important as new strains of the virus may evolve, or future coronaviruses may emerge, that are resistant to current vaccines. The application of transcriptomic RNA sequencing of infected samples may shed new light on the pathways involved in viral mechanisms and host responses. We describe the application of “dual RNA-seq” analysis to consider both the host and pathogen transcriptomes simultaneously, to investigate for the first time the *co*-regulation of human and SARS-CoV-2 genes. Together with differential expression analysis, we describe the tissue specificity of SARS-CoV-2 expression, an inferred lipopolysaccharide response, and co-regulation of *CXCL’s, SPRR’s, S100’s* with SARS-CoV-2 expression. Lipopolysaccharide response pathways in particular offer promise for future therapeutic research and the prospect of subgrouping patients based on chemokine expression that may help explain the vastly different reactions patients have to infection. Taken together these findings illuminate previously unappreciated SARS-CoV-2 expression signatures, identify new therapeutic considerations, and contribute a pipeline for studying multi-transcriptome systems.

## Introduction

Currently, the SARS-CoV-2 pandemic, which is caused by the coronavirus disease 2019 (COVID-19), has a global mortality rate that is still unknown ^1,2^. Its first appearance was reported in December 2019 and has since spread to 213 countries and territories and has caused over 100 million confirmed cases ^1,3–5^. The SARS-CoV-2 infection has been reported to cause a variety of symptoms including fever, cough, fatigue, shortness of breath, and abnormalities in the chest as determined by CT ^6–8^. Severe cases manifest with acute respiratory distress syndrome and lung injury, leading to morbidity due to damage to the alveolar lumen leading to inflammation and pneumonia ^9,10^.

A variety of model systems and tissue types have been used to study the transcriptional response to SARS-CoV-2 infection in an effort to better understand the molecular basis of COVID-19 ^11,12^. These studies have revealed a cytokine, chemokine, and immune response to SARS-CoV-2 infection. This gene signature has been useful in understanding the biology of the COVID-19 disease. The differential expression of these gene families is, therefore, a consistent theme for the SARS-CoV-2 infection, but what hasn’t been deeply explored is how SARS-CoV-2 genes associate with and activate a host response in human gene expression.

We developed an unbiased multi-transcriptome read alignment method to investigate the transcriptomes of human and virus together in the same sample. Previous studies analyzed only the human transcriptome, which provided important information about what genes and pathways are *differentially expressed* between infected compared to uninfected samples. However, as illustrated by countless gene expression analyses from microarrays to RNA-seq, a complementary systems-level view can be achieved by looking at the *co-expression* of genes that may pick up on more subtle, but still significant associations missed by differential expression. Our method leverages *dual-RNAseq* to quantify transcripts from both the host and pathogen together, which has shown promise in other systems ^13,14^. Dual-RNAseq originally required additional library enrichment to detect rare classes of transcripts. However, modern sequencing now yields a high enough read depth to provide accurate quantitation of the entire host and pathogen transcriptomes without the need for additional library enrichment steps.

We investigated the utility of dual-RNAseq to study samples infected with the SARS-CoV-2 virus that makes it possible to correlate human genes with specific viral genes. We applied the analysis to both cell lines and patient samples and used multiple correlative methods including average linkage dendritic clustering, Pearson correlation networks, and Pagerank network importance. We derive a consensus network implicated by these multiple views that sheds new light on the roles of human genes and pathways in SARS-CoV-2 infection.

## Results

We built a dual RNA-seq analysis pipeline (**dRAP**) to map all transcripts of infected cells to either the host or viral genomes in an unbiased manner (Fig.1A; see Methods). Our hypothesis is that quantification of both host and viral transcripts would enable a more sensitive and specific association of host pathways responsive to SARS-COV-2 (SARS-CoV-2) infection. While many publications report results based on the expression of human genes in SARS-CoV-2 infected cells, to our knowledge our attempt represents the first to estimate both host and virus transcription simultaneously in the same sample for SARS-CoV-2.

### Robust viral transcript detection in human cell lines

As a preliminary test, we first applied dRAP to the analysis of human cell lines for which the amount of infected virus was experimentally controlled (Fig. 1B; Figure 2A). Blanco-Melo et al. ^11^ exposed a variety of human respiratory cell lines with SARS-CoV-2 virus. In that study, the authors found a robust host transcriptional response to infection for an ACE2 receptor-enhanced alveolar basal epithelial cell line (A549) as well as a bronchial epithelial cell line (NHBE). As the A549 and NHBE cell lines show robust host response and have established viral levels, we reanalyzed the data with dRAP to jointly analyze both the host and viral transcriptomes.

**Figure 1.**
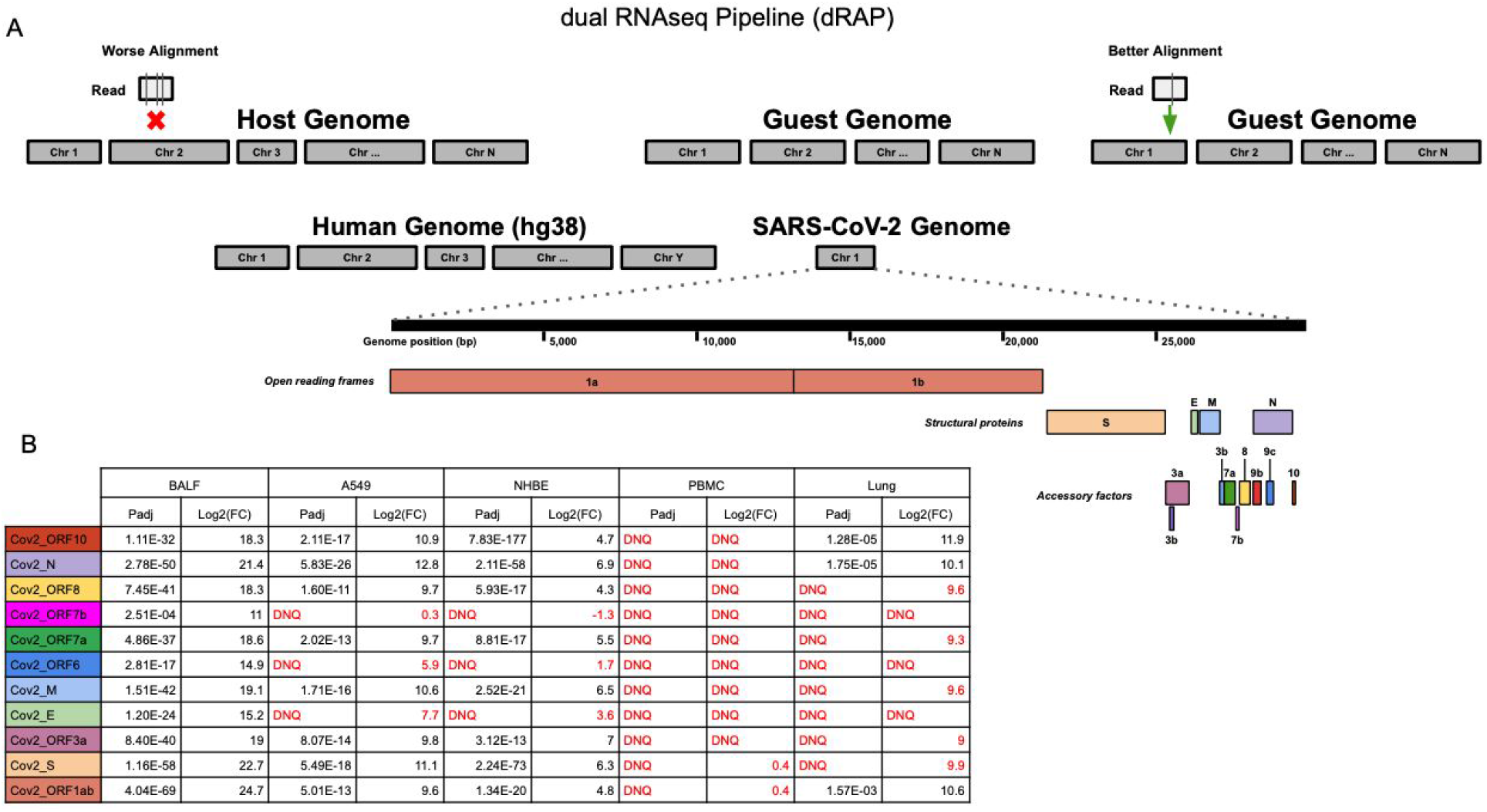
(A) The dRAP pipeline follows a dual RNAseq methodology by creating a single reference index for RNAseq read alignment with STAR by appending “guest” genomes to the host genome. This allows for simultaneous, unbiased alignment of reads with multiple transcriptomes where the best overall alignment is selected. For example, aligning a single read would result in 3 mismatches (vertical lines in box under “Worse Alignment”) if aligned to the host genome (red “x” marks host alignment location) compared to 1 mismatch (vertical line in box under “Better Alignment”) aligning to the guest genome (green arrow). Both the human host genome (hg38) and the SARS-CoV-2 guest genome (NC_045512v2) are included, which has a set of genes that are designated as open reading frames, structural proteins, or accessory factors. (B) SARS-CoV-2 differential gene expression results for infected patient tissue and cell line samples compared with non-infected samples. Patient tissue types show dramatically different expression profiles with PBMC and Lung biopsy tissue rarely ever passing detection limits while BALF tissues show robust expression in infected patients. Cell lines also display strong SARS-CoV-2 expression although the magnitude of fold change was far less than that observed in BALF samples. “DNQ” stands for “did not qualify”, which indicates genes that failed Cook’s distance filtering in DESeq2 analysis.

**Figure 2.**
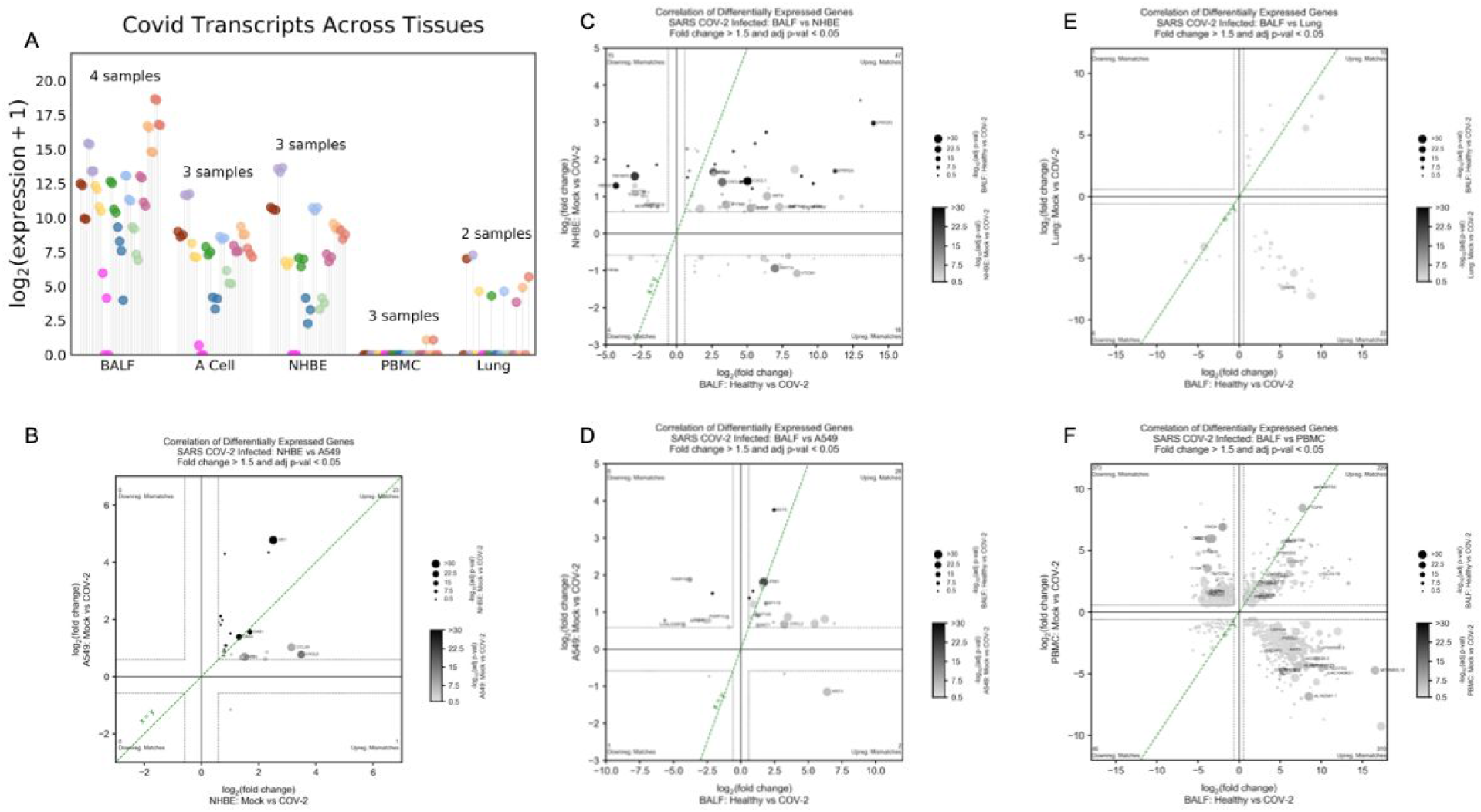
(A) dRAP is sensitive enough to detect subtle differences in SARS-CoV-2 transcripts quantities resulting in differential expression within the SARS-CoV-2 transcriptome. SARS-CoV-2 expression is also shown to be highly dependent on the system being studied. Patient BALF samples show high amounts of SARS-CoV-2, while PBMC and Lung patient samples display low or no SARS-CoV-2. (B-F) Log2 fold change comparison between differential expressions in infected samples against non-infected samples shows that the tissue specificity of SARS-CoV-2 extends to the degree of concordance observed in the human transcriptome. Statistically significant concordance was observed between NHBE and A549 cell lines (p<1e-5)(B) and BALF patient samples with NHBE (p<0.05)(C) and A549 (p<0.01)(E) cell lines using the chisquare test. However, there was no concordance observed in BALF versus Lung (p=0.26)(D) and a significant discordance versus PBMC (p<1e-38)(F) patient samples. This suggests that the lack of SARS-CoV-2 expression observed in Lung and PBMC samples is also associated with significantly altered human expression, making these tissue types not ideal for learning SARS-CoV-2 mechanisms.

Importantly, dRAP provides a quantification of the amount of viral transcript in a host cell, which could be leveraged for downstream correlation-based analyses to implicate pathways of host response. Thus, while there is high concordance of the particular transcripts detected, dRAP reports a range of fold changes, with more dramatic overexpression detected in the A549 cell line compared to NHBE, potentially due to contamination of the mock-treated NHBE cells with SARS-CoV-2. Whether the differences are due to technical artifact or biological factors, these observations support the idea that dRAP’s quantifiable differences can serve as the basis for studying regulation dynamics associated with infection by jointly analyzing the viral and host transcripts together.

The relative statistical significance of genes was also found to be consistent between the cell lines (Kendall rank correlation 0.69, p<0.01). For example, the genes core to the infection of the virus --- *ORF10, S, N*, and *M* -- were found to be the highest expressed genes in both A549 and NHBE. Out of the eleven SARS-CoV-2 transcripts, eight were detected consistently in both cell lines, suggesting viral transcripts can be determined robustly by dRAP in multiple host cell types. In addition, there is significant concordance of differentially expressed human genes between SARS-CoV-2 and RSV infections (Chi-Square 45.1, p<1e-10) consistent with the report by the original authors ^11^ (Suppl. Figure 1).

### Variable detection of viral transcripts in human tissues

We next applied dRAP to the estimation of viral transcription in human tissues from patients with known infections. The differences in viral quantities across patient samples may lead to differences in host response. Therefore, correlating human transcription changes that track closely with particular changes in viral transcripts might then identify host responses that are more likely to be directly associated with the virus compared to the many potential indirect effects that have been found through differential expression analysis.

To this end, we first searched for human cells that express the most consistent levels of SARS-CoV-2 virus. We expected that lung tissue would show the highest amount of viral transcription since previous reports that the virus invades alveolar epithelial cells ^15^. However, to our surprise, the most striking finding was that many SARS-CoV-2-infected patient samples from the lung or blood showed no SARS-CoV-2 expression at all. Specifically, PBMC samples had one patient that showed a normalized transcript count of 1.1 for *ORF1ab* and *S*, while all other samples and all SARS-CoV-2 genes had transcript counts of 0 (Fig. 1B; Figure 2A). No SARS-CoV-2 gene passed DESeq2 Cook’s distance filtering for the PBMC samples. Similarly, lung biopsy samples showed inconsistent SARS-CoV-2 expression (Fig. 1B; Figure 2A). One patient showed SARS-CoV-2 expression while the other patient had none. However, even for the patient that did display SARS-CoV-2 gene expression, the expression levels were far less robust than that observed in cell lines.

In stark contrast with these other patient samples, SARS-CoV-2 patient samples from Bronchoalveolar lavage fluid (BALF) showed very robust SARS-CoV-2 expression that even exceeded the levels observed in infected cell lines (Fig. 1B; Figure 2A). The most significant SARS-CoV-2 gene was *ORF1ab*, followed by the *S, N*, and *M* genes. The overall profile of BALF samples has a few notable differences from that of the infected cell lines, including a much more significant overexpression of the SARS-CoV-2 *ORF1ab* gene and that the *E, ORF6*, and *ORF7b* genes were also significantly overexpressed in BALF samples but not in cell lines. Outside of these differences the cell lines show similar features to the BALF SARS-CoV-2 profile at a lower expression level, including the predominant overexpression of the *S, N*, and *M* genes. Overall, the relative significance of SARS-CoV-2 genes were highly concordant between BALF and A549 (Kendall rank correlation 0.49, p<0.05) and NHBE (Kendall rank correlation 0.56, p<0.05) samples. In contrast, BALF and PBMC (unable to compute Kendall rank correlation because all transcripts were undetected in PBMC samples) and Lung (Kendall rank correlation 0.23, p=0.37) were shown to be discordant. Because of the similarities between BALF patient samples and cell lines and the absence of a robust SARS-CoV-2 expression profile in PBMC and lung biopsy samples, we used BALF, NHBE, and A549 samples in the following host-virus joint analysis to implicate host pathways associated with SARS-CoV-2 infection.

### Concordant expression changes in BALF and cell lines

Identifying differentially expressed (**DE**) genes between two conditions can be a powerful way to uncover regulatory mechanisms implied by the transcriptional changes. Because expression can reflect many cell types and processes occurring in parallel, they reveal both direct and indirect effects. Given that the virus is detected in BALF and not PBMCs and lung, we reasoned that host expression changes in BALF would reflect a higher degree of direct responses to infection, compared to lung and PBMCs. To test this, we asked if the BALF DE genes were more comparable to the cell line DE genes relative to the DE genes derived from PBMCs and lung.

The total expression profile of these samples were then compared against each other to determine common and diverging signatures in the various patient tissue samples and cell lines. First, the profile of the NHBE and A549 cell lines were compared (Figure 2B). Overall, there were 23 matching directional expression changes and 1 mismatched directional expression change in genes meeting Fold Change and p-value thresholds for both NHBE or A549 cells, demonstrating a significant concordant relationship (Chi-Square 20.2, p<1e-5). Genes showed a common up-regulation between the two cell lines, which is clearly observed in the plot through changes in size and color indicating increasing significance. Some of the most significant changes were observed in genes with roles in antiviral response (MX1, IFI27, IRF9, OAS1, OAS3), and chemokine signaling (CXCL5).

Comparing BALF patient samples with NHBE cells there were 51 matching directional expression changes and 33 mismatched directional expression changes (Figure 2C), demonstrating significant concordance (Chi-Square 3.9, p<0.05). Most genes show common up-regulation (47 genes), including *SPRR2D, SPRR2A, PLAT, CXCL1*, and *CXCL2*. The comparison of BALF with A549 cells produced a similar result (Chi-Square 9.3, p<0.01)(Figure 2D) with 29 matching directional expression changes and 10 mismatched directional expression changes.

Next, we compared BALF with lung and PBMC samples. BALF and lung samples were found to have 16 matched directional expression changes and 23 mismatched directional expression changes, demonstrating that the majority of gene expression changes are discordant, but without reaching statistical significance(Chi-Square 1.3, p=0.26)(Figure 2E). In a more extreme manner, BALF and PBMC samples showed 275 matched directional expression changes and 683 mismatched directional expression changes, demonstrating that the majority of gene expression changes are very significantly discordant (Chi-Square 173.7, p<1e-38) with BALF samples (Figure 2F). Therefore, for both PBMC and lung samples there was a dominant discordance in gene expression changes compared to BALF. This is consistent with our observations above that find robust SARS-CoV-2 expression levels in BALF compared with undetectable levels in PBMC and lung.

### Multi-view coexpression reveals a consensus human transcriptional network

The estimates of viral and host RNAs for the same samples provided by dRAP enable investigating host regulatory pathways through a coexpression (CE) analysis to identify human transcripts most correlated with viral transcripts. CE could complement DE to find human genes directly associated with viral infection by detecting more subtle patterns of transcripts fluctuating concordantly with particular viral products that may suggest regulatory connections between the viral and host genes. The BALF human tissue samples and the two cell lines were used for CE as the above analyses found these samples provided robust viral RNA expression.

To prioritize candidate genes and pathways, we elected to use three different CE strategies to see if any genes and pathways were nominated by one (or more) viewpoint(s). First, we collected the top *k* most correlated human transcripts using Pearson correlation for each viral protein and saved the union of these human proteins. Next, we performed a hierarchical clustering analysis on the Pearson correlation matrix to determine if there existed a clade, or clades, enriched for viral proteins. Third, we performed a PageRank analysis on the Spearman correlation matrix to determine which host genes are central to the host-virus correlation network. Candidate host genes were collected from each of these approaches, compared to each other as well as to the list of genes from DE analysis.

A second approach we used was to take the entirety of normalized transcript counts passing DESeq2 Cook’s distance filtering and cluster their expression using an average linkage distance between pairwise Pearson correlations to create dendrograms with various clades. BALF samples and NHBE and A549 cell lines infected with SARS-CoV-2 all had extremely localized co-expression of SARS-CoV-2 genes with each other. This is evidenced by the fact that in NHBE cells (Figure 3A-B) and BALF samples (Suppl. Figure 2A) all SARS-CoV-2 genes passing Cook’s distance were co-expressed in the same clade as each other. A549 cells were similar with the exception of the SARS-CoV-2 *N* gene, which was located in an adjacent clade (Suppl. Figure 2B). The CE-dendro analysis clustered some genes with SARS-COV-2 genes (Suppl. Figure 3) that were not implicated by differential expression analysis as they failed to pass Benjamini-Hochberg corrected significance tests. These include *MYC, NFKBIA*, and *DDX1* that are implicated in proto-oncogenic pathways ^16^, immune response in lungs ^17^, and host cofactor enhancement of SARS-CoV-1 replication ^18^, and thus of possible relevance to SARS-COV-2 mechanisms.

**Figure 3.**
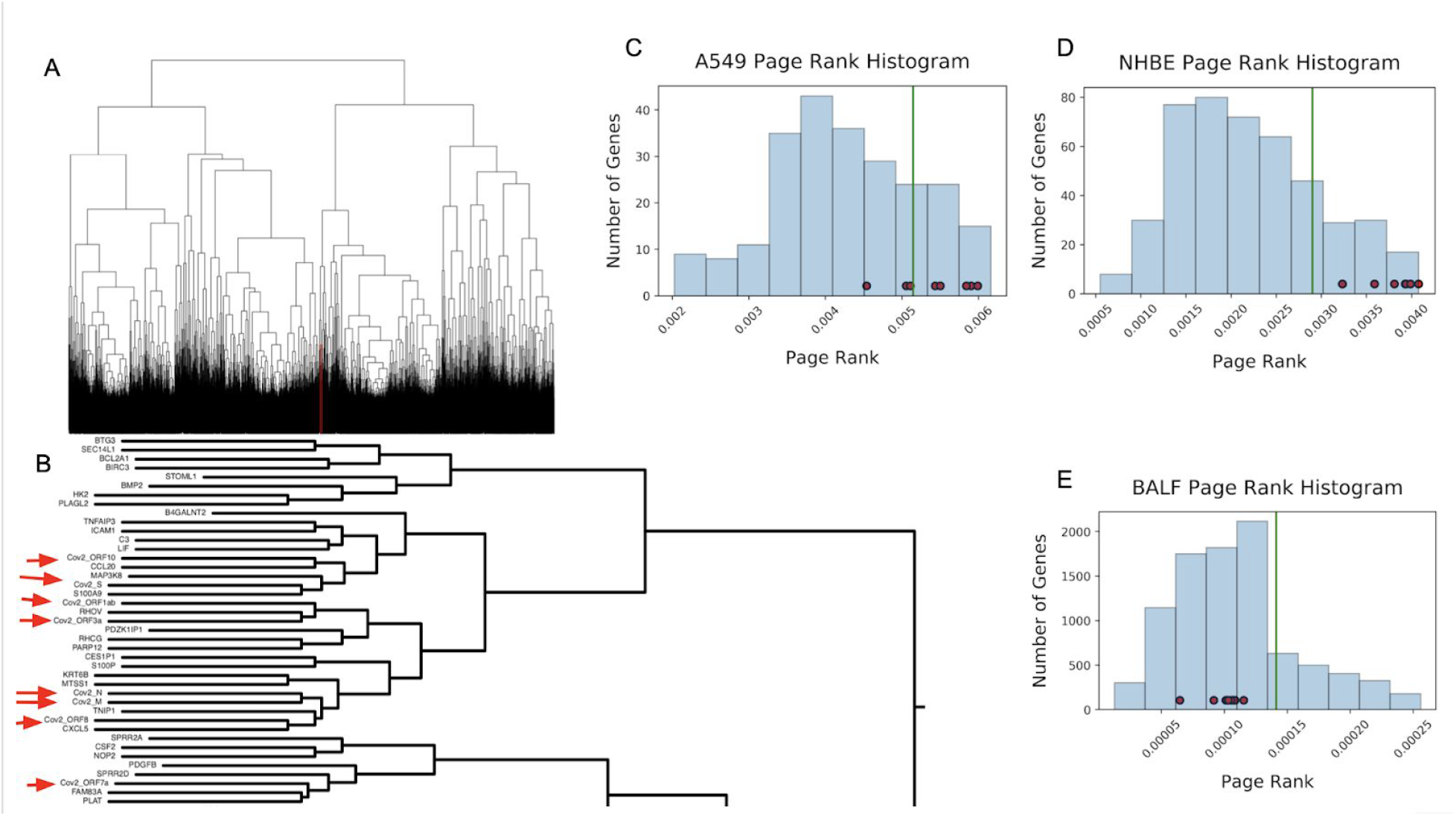
By clustering expression SARS-CoV-2 and human expression patterns concurrently we observed that SARS-CoV-2 transcripts were localized in a small clade visualized in red (A). This clade of coexpression with SARS-CoV-2 transcripts contains a set of genes associated with SARS-CoV-2 mechanisms in infection (B). The histogram distribution of PageRank values for the A549 (C) and NHBE (D) cell lines shows that the SARS-CoV-2 genes are highly influential. However, in BALF samples (E), SARS-CoV-2 genes are at the lower end of the PageRank distribution likely due to the numerous differentially expressed genes creating a much larger set than that for the cell lines. In A-C, the green line marks the 80th percentile in the distribution and the small red nodes along the distribution represent SARS-CoV-2 genes.

The third view used genes with significant differential expression (padj<0.05) to create a network of genes with an *R*^2^ > 0.98 with a SARS-CoV-2 gene (see Methods). The NHBE and A549 networks have a very similar architecture, whereas the BALF network displays a much denser network of expression (Suppl. Figures 4-6). All three of these networks display very similar gene signatures of Chemokines, *SPRR’s, S100’s*, viral response, and interferon response genes. The CE-Net for the strong linear relationships found between SARS-CoV-2 genes and human genes displays a very modularized and sparse architecture of particular gene groups being localized and indicated in a group of pathways (Suppl. Figure 7). This is a desirable architecture to visually parse and extract gene subgroups that coordinate to elicit specific responses. A few examples of important gene subgroups in Suppl. Figure 7 include the gene modules indicative of a lipopolysaccharide response and chemokine/cytokine activity (*CXCL5, CXCL8, CCL20, HIF1A*), cornification and epithelial cell differentiation (*SPRR2A, SPRR2D, SPRR2E, PI3, KRT6B, ESF1, RHCG, MTSS1*),and antiviral response (*OAS1, MX1* and *PARP9, DTX3L*).

**Figure 4.**
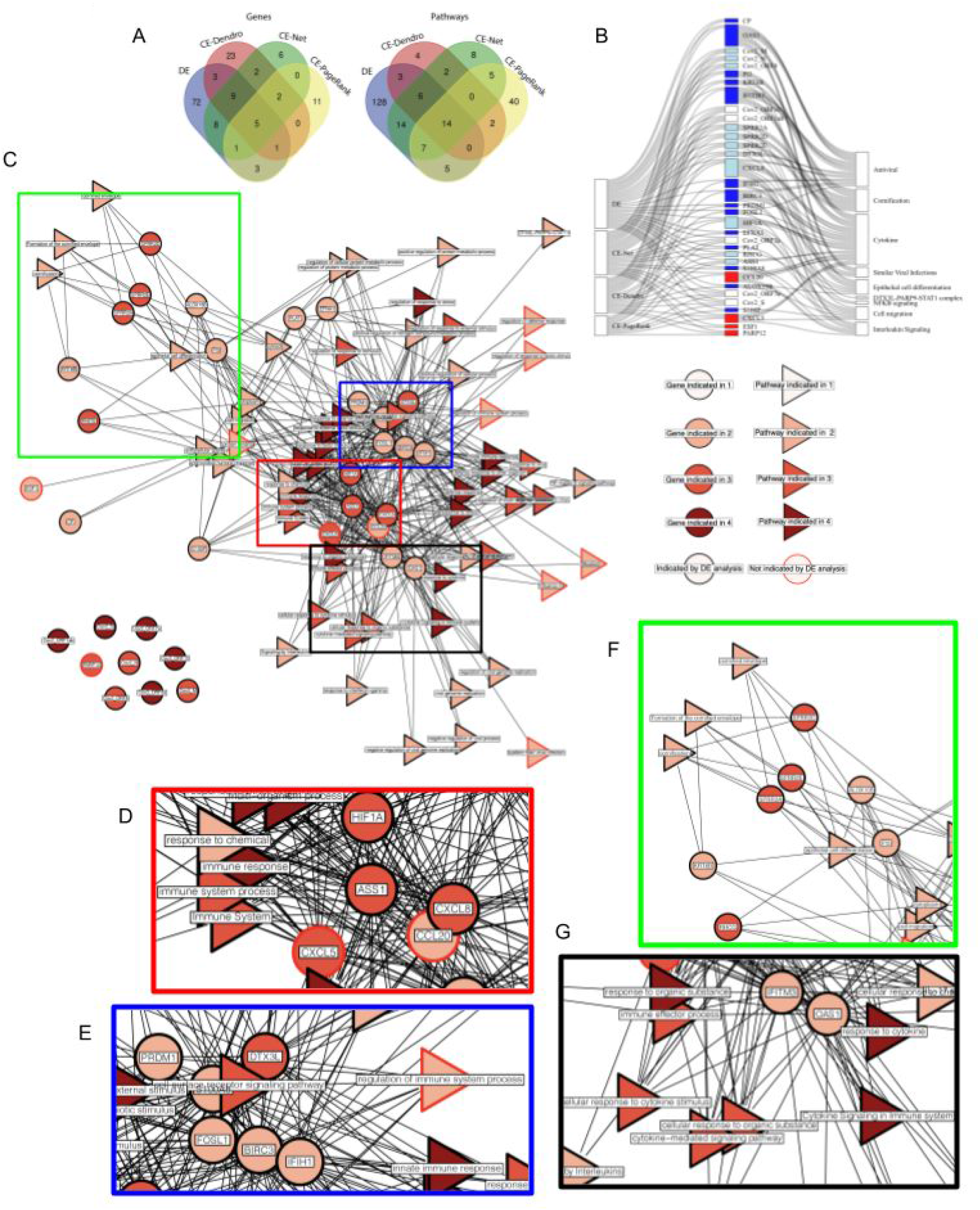
(A) The overlap in gene and pathway results indicated by DE, CE-Dendro, CE-Net, and CE-PageRank views by venn diagram. (B) Sankey diagram showing genes found by at least two of four views and pathway themes in which they participate. Genes are colored based on the views that they are found in, white, light blue, and dark blue indicate that DE and 1, 2, or 3 other CE views, respectively, and red for a gene that is not found by DE, but was by a CE view. Genes are connected to themes created from a list of gProfiler pathways (Suppl. Table 1). (C) The consensus network for the cross-analysis overlaps between these four views results in four gene modules. Genes are displayed as circular nodes and pathways are displayed as triangular nodes. Nodes with red borders represent results that did not manifest in traditional differential expression analysis, and, therefore, demonstrate the power of the dRAP pipeline to not only highlight important findings in DE analysis, but to also reveal previously hidden signatures. (D) The red module (*CXCL5, CXCL8, CCL20, ASS1, HIF1A*) is indicative of chemokine activity, cytokines, and a lipopolysaccharide response. (E) The blue module (*IFIH1, PRDM1, BIRC3, FOSL1, DTX3L, S100A8*) indicates an innate immune response. (F) The lime module (*SPRR2A, SPRR2D, SPRR2E, KRT6B, ALOX15B, PI3*) indicates cornification/keratinization and epithelial cell differentiation changes. (G) The black module (*OAS1, IFITM3*) indicates regulation of viral genome replication.

The final approach used weighted PageRank to find genes with high influence among a set of significant DE genes. In A549 cells, many SARS-CoV-2 genes (*S, ORF3a, ORF1ab, ORF7a, ORF10*) were indicated as above the 80th percentile of influence amongst significant DE genes (Figure 3C). Similarly, SARS-CoV-2 genes (*M, S, N, ORF8, ORF10, ORF3a, ORF7a, ORF1ab*) were largely indicated as being above the 80th percentile of influence (Figure 3D). However, in BALF patient samples, no SARS-CoV-2 were above the 80th percentile of influence (Figure 3E). One likely reason for this would be that the number of DE genes is over 20 times more numerous than either of the cell lines. Therefore, the gene regulatory network driving the conditional response is hidden behind more noise, likely due to the inherent increase in variability between patient samples compared with cell lines. This suggests that many of the DE genes in BALF samples may be more indicative of patient variability than of a response to SARS-CoV-2 infection. CE for weighted PageRank shows a similarly sparse and modular architecture to that observed by CE-Net (Suppl. Figure 8). However, the subgroups participating had some notable differences in the participating modules, which consisted of the orange module (*S100P, PROS1, PTAFR*), pink module (*ICAM1, HBEGF, INHBA*), purple module (*CXCL5, ASS1, DTX3L, BIRC3, IFIH1*), and black module (*MAFF, HDGF, CSNK1E, SAMD9, ITGA5, MCFD2*). Finally, the CE-PageRank network (Suppl. Figure 8) shows a variety of unique, modularized results including purple module (*CXCL5, ASS1, DTX3L, BIRC3, IFIH1*)(which was similar to the red and blue modules observed in the consensus network (Figure 4D-E)), pink module (*ICAM1, INHBA, HBEGF*), orange module (*PROS1, PTAFR, S100P*), and black module (*MAFF, HDGF, CSNK1E, SAMD9, ITGA5, MCFD2*). Genes of potential import to highly include *ICAM1*, which has been implicated in viral entry and survival ^19^, *S100P*, implicated in calcium binding and human papillomavirus (HPV) ^20^, and *SAMD9*, shown to have antiviral properties ^21^.

Relative to DE, the CE views produce some common, but primarily distinct, genes and pathways (Figure 4A). We collected 102 genes from the DE analysis that were implicated by two out of the three separate DE analyses run on BALF, NHBE or A549 (see Methods; Suppl. Figure 9). The pathways in the core of the DE results were highly consistent and reminiscent of those described previously by the original authors ^11^. We found that the candidate genes suggested by the three different CE approaches are distinct from the DE results (overlaps of 18, 23, and 10 with the CE-Dendro, CE-Nets, and CE-PageRank results, respectively). Of the 102 DE genes, a majority (72 genes) were only indicated through DE, suggesting that DE produces a large number of results that are exclusively a statistically significant conditional response and are not indicative of linear correlation to the source (CE-Nets), co-regulation with the source (CE-Dendrogram), or the most influential players in the geneset (CE-PageRank).

To reveal a core host regulatory response associated with SARS-COV2 infection, we constructed a consensus from the results of DE, intersected with any of the three CE views and plotted the results as a Sankey diagram (Figure 4B). Thematic biological processes were identified using the gProfiler tool ^22^ on the resulting consensus to implicate pathways recorded in the gProfiler ontology database that are statistically enriched among the gene set (see Methods). SARS-COV-2 genes themselves are not included in the consensus as gProfiler lacks annotations for this virus. The resulting consensus shares much in common with the CE-Net, which is expected given the common results revealed by the views as discussed above. Several genes and pathways were commonly implicated by DE and the CE views (e.g. 30 genes by DE and at least one CE; 16 genes by DE and two or more CEs). Five genes, all viral encoded

--- *Cov2_ORF7a, Cov2_ORF1ab, Cov2_S, Cov2_ORF10*, and *Cov2_ORF3a*, which were first quantified using dRAP -- and 14 pathways indicative of a Immune and Defense response were found by all four views. Two genes -- *CXCL5*, a chemokine, and *PARP12*, an interferon-stimulated gene involved in regulating inflammation -- were found by the three CE views, but not by DE. The nine genes (*Cov2_ORF8, Cov2_M, Cov2_N, SPRR2A, SPRR2D, SPRR2E, RHCG, HIF1A, CXCL8*) found by DE, CE-Nets, and CE-Dendrogram, but not CE-PageRank could also be of interest for tight association with the virus albeit not central to the known pathway membership interconnections that influence the PageRank analysis.

Viewing the consensus as a network (Figure 4C) highlights several sub-modules of densely interconnected genes and pathways (colored boxes, Figure 4D-G). The first module (red box: *CXCL5, CXCL8, CCL20, ASS1, HIF1A*; Figure 4D) consisted of genes belonging to several pathways expected to be implicated including Chemokine activity, IL-17 signaling pathway, viral protein interaction with cytokine and cytokine receptor, cellular response to lipopolysaccharide, and the ‘ASATCAAAG’ TCF-3 motif. The module is supported by previous findings that SARS-CoV-1 infection stimulates the lipopolysaccharide receptor, TLR4, shown to produce an immune response ^23^ and lead to disease pathogenesis ^24^. The second module (blue box: *DTX3L, IFIH1, BIRC3, S100A8, PRDM1, FOSL1*; Figure 4E) implicates roles for the innate immune system’s defense response. *DTX3L* and *PARP* may be of particular interest based on previous findings that they are required for an interferon response to certain coronaviruses ^25^ and *PARP12/14* are required to inhibit the replication of the macrodomain (a subunit of the transmembrane viral protein, nsp3) of coronaviruses and to produce the optimal IFN response ^26^. The third module (lime box: *SPRR2A, SPRR2D, SPRR2E, KRT6B, ALOX15B, PI3, RHCG*; Figure 4F) implicates the involvement of programmed cell death, epithelial cell differentiation and cornification and antiviral response through a *DTX3L-PARP* axis. This gene subgroup is characterized by Keratinization, Keratinocyte cell differentiation, cornification, formation of the cornified envelope, Epithelial cell differentiation, Epidermal differentiation, programmed cell death, and structural constituents of skin epidermis. Finally, a fourth module (black box: *OAS1, IFITM3; Figure 4G*) indicates pathways of viral genome replication and cellular response to type 1 interferon. In summary, the consensus network reveals many expected aspects of response involving chemokines and inflammation but also some potentially new processes such as cornification that may implicate a specific apoptotic mechanism involving particular host cells (e.g. keratinocytes).

## Discussion

Using a dual RNA-seq analysis pipeline (dRAP) to detect both host and pathogen transcriptomic gene expression, we implicate genes involved in viral infection and response using both differential expression (DE) and coexpression (CE) analyses. To elucidate therapeutic implications, we applied dRAP to SARS-CoV-2-infected patient samples and cell lines. For the first time to our knowledge, we quantify the levels of SARS-CoV-2 transcripts in human patients and cell lines. This new outlook revealed that the most strongly and consistently expressed transcripts were those that play essential roles in viral survival and propagation. The *S, N*, and *M* genes -- which are key to viral replication (*N* gene), assembly (M gene), release (*M* gene), attachment (S gene), and entry (S gene)^27,28^ -- had the highest levels of statistically significant expression.

dRAP suggested an appreciable difference in viral transcript expression between patient tissue types and found that BALF had the most robust levels. PBMCs exhibited low (or zero) levels of SARS-CoV-2 transcript expression, with no transcript detected as differentially expressed between control and infected conditions (DESeq2 using Cook’s distance filtering; Figure 1B, Figure 2A). Similarly, lung biopsies lacked robust expression of SARS-CoV-2 transcripts with all but one sample giving detectable levels (Figure 1B, Figure 2A). In contrast, BALF samples were found to express every SARS-CoV-2 transcript at extremely high levels (at least 14 times higher in infected samples compared to controls; Figure 1B, Figure 2A).

In addition to the robust viral response, the *human* transcriptional response of BALF also matched more closely with SARS-CoV-2-infected cell lines than the other tissues matched to cell lines. dRAP identified that the NHBE cell line produced the highest magnitude of viral transcript expression (magnitudes higher than the A549 cell line, which also had some detectable viral transcript levels). The human transcriptional response, as measured by differential expression (DE) was found to be most similar between BALF and NHBE (Figure 2C) with the other tissues having little in common with cell lines or BALF (Figure 2E-F). Interestingly, the commonality between BALF and the NHBE cell line was even greater than when the cell lines were compared to each other (Figure 2B). These findings both underscore the relevance of using BALF as the tissue to quantify the gene expression programs of SARS-CoV-2 infection while also affirming that some cell lines (e.g. NHBE) may offer approximate surrogate laboratory models.

By providing both virus and human transcript estimates in each sample, dRAP enables the use of coexpression (CE) in addition to DE. We investigated three different views to explore the roles of human genes and pathways relevant to viral infection implicated by CE. DE measures the statistically significant conditional response to infection (Suppl. Figure 9) that may nominate genes with either direct or indirect viral associations resulting in many genes and pathways relative to CE (off-white nodes), with little to no co-regulation with SARS-CoV-2 genes (CE-Dendro or CE-Nets), or central to its correlation network (CE-PageRank). The consensus of the DE and CE views (Figure 4A, SFig.10) provide a coherent overview of SARS-COV-2 infection and host response supported by literature observations of previous coronaviruses, viral response machinery, and symptoms observed in COVID-19 patients.

The modules of the consensus network suggest that SARS-CoV-2 may initiate a response via lipopolysaccharide through increases in chemokine and cytokine activity, triggering an influx of intracellular calcium that induces the migration and programmed death of epithelial cells through a hardening of the membrane by a keratinization/cornification process ^23^. These findings are supported by previous observations that chemokines activate a response in multiple cell types and tissues (including epithelial and leukocyte) to produce high intracellular calcium and a migratory phenotype ^29–32^. Imbalances and compensation among chemokines may predict response to infection ^33^. Indeed, mutations in chemokine-associated genes have been associated with severe cases ^34^. Chemokine pathways may underlie symptoms that coincide with the various observations of COVID-19 symptoms including a respiratory mucosal immune response ^33^, inflammatory bowel disease ^35^, interstitial lung disease ^36^, asthma ^37^, and Eosinophilic Pneumonia (EP)^38^, which is associated with “progressive shortness of breath (dyspnea) of rapid onset and possibly acute respiratory failure, cough, fatigue, night sweats, fever, and unintended weight loss.” Understanding the full scope of possible chemokine pathway engagement and how to inform therapeutic approaches is of great interest.

In summary, this work suggests that specific human patient tissues are implicated in robust SARS-CoV-2 expression, namely, Bronchoalveolar lavage fluid (BALF). NHBE and A549 infected cell lines offer a faithful representation of the SARS-CoV-2 induced gene expression signature when compared to BALF, but PBMC and lung biopsy patient samples do not show robust SARS-CoV-2 expression or capture the appropriate representation of the conditional human gene expression response. Additionally, the coexpression analysis enabled by dRAP produced a coherent hypothesis that predicted a possible mechanism by which COVID-19 may progress from the initial SARS-CoV-2 infection to patient symptoms. Taken together, these findings shed light on the molecular pathways implicated in COVID-19, a worldwide health crisis that desperately needs therapeutic targets, demonstrating the power of the joint host-virus view provided by dual RNA-seq to generate specific hypotheses from human transcriptomic expression changes.

## Methods

### Dual RNA-seq to simultaneously map virus-host transcriptomes

In order to quantify the host and pathogen transcriptomes, we implemented a dual RNA-seq ^13,14^ analysis pipeline (dRAP). dRAP takes a series of reference FASTA files and their corresponding GTFs and concatenates them into a single FASTA and GTF, which is subsequently used to create a mapping index. The human reference FASTA and ENSEMBL GTF for hg38 and the SARS-CoV-2 reference FASTA (NC_045512v2) and REFSeq GTF were downloaded from the UCSC Genome Browser ^39^. STAR was then used with the parameters runMode, sjdbOverhang, and genomeSAindexNbases as ‘genomeGenerate’, 100, and 6, respectively, to create a merged hg38/NC_045512v2 index ^40^. Following the creation of the merged index, each sample was then mapped to the index using STAR to get transcription counts with the parameters outSAMtype, twopassMode, outFilterMultimapNmax, and quantMode set as ‘BAM SortedByCoordinate’, ‘Basic’, 1, and ‘GeneCounts’, respectively. This generated a ‘ReadsPerGene’ for each sample, which was then used as input for DESeq2 count normalization and differential gene expression (DGE) determination ^41^. Gene expression levels, as read counts, were estimated and filtered by Cook’s distance and nominal p-values were corrected for False Discovery Rates (FDR) and a significance threshold was set at FDR < 0.05.

### Differential Expression (DE) Analysis

To test whether dRAP is capable of detecting and quantifying host and pathogen transcripts, we used two previously published datasets on patients and cell lines infected with SARS-CoV-2 ^11,12^. NHBE and A549 cell lines (Wild-type n=3 and SARS-Cov-2 infected n=3 each) and postmortem Lung Biopsies (Healthy n=2 and SARS-CoV-2 infected n=2) were processed from the Tenoever data. Healthy (n=3) and SARS-CoV-2 infected (n=3) PBMC’s and healthy (n=3) and SARS-CoV-2 infected (n=4) BALF patient samples were processed from the Chen data. DESeq2 DGE of SARS-CoV-2 transcripts was determined for each tissue. Kendall rank correlations were calculated for the padj values of SARS-CoV-2 transcripts between each tissue type. Genes that did not qualify for statistics based on the DESeq2 cook’s distance criteria were included as padj of 1, i.e. as the last rank in the set.

### Comparing Human Differential Expression between samples and cell lines

Because of the differences in SARS-CoV-2 transcripts detected between tissue and sample types, we wanted to determine if these differences also were reflected by the host transcriptional response. A one-to-one comparison of the log2 Fold-Change for each tissue against each cell line was plotted to determine common differential gene expression patterns. Each panel displays the log2 Fold-Change of one group versus the log2 Fold-Change of another by plotting genes that exceed significance and fold change thresholds in each group. Each plotted gene is sized according to its significance (-log10 adjusted P-value) for the x-axis group and colored according to its significance for the y-axis group. Genes that were up-regulated and met threshold criteria for both groups were recorded as up-regulated matches. Figures display genes that met the threshold criteria of |Fold-Change| > 1.5 and padj < 0.05. A Chi-square test was then used to determine the likelihood of the distribution of genes with matched and mismatched expression directionality between the two groups. These DGE were then used as input for the gProfiler:GOSt functional profiling ^22^. Common genes and ontologies in SARS-CoV-2 infected groups were recorded for use in downstream analysis.

### Co-Expression (CE) Analysis

Because the differences in SARS-CoV-2 expression appeared to result in notable changes in host transcriptome response, we hypothesized that gene co-expression with SARS-CoV-2 may illuminate mechanisms of action and therapeutic targets. To this end, we utilized three different strategies as part of a Co-Expression (CE) analysis. These included analyses of genes demonstrating high correlation with SARS-CoV-2 (CE-Net), genes clustered with SARS-CoV-2, and genes with high weighted-PageRank influence (CE-PageRank).

For groups infected with SARS-CoV-2, genes passing Cook’s distance filtering with DESeq2 were clustered using average linkage with the R package ‘hclust’ ^42^. The resulting dendritic tree was then cut into 200 clades of co-expression. Clades containing SARS-CoV-2 transcripts were then further subdivided into 5 clades. After this final subdivision, co-regulated genes participating in clades containing SARS-CoV-2 transcripts were used as input for gProfiler:GOSt functional profiling ^22^. The input genes and resulting pathways were separated and used in downstream analysis.

DE genes with benjamini-hochberg corrected p-values less than 0.05 were subsetted for PageRank analysis. We used the topological overlap matrix (TOM)^43^ generated from a Pearson correlation matrix with a soft thresholding parameter of 30 to create a weighted gene network using pairwise complete observations and then ran weighted PageRanks with a damping parameter of 0.9 on each of the 5 sample groups ^44^. Genes with PageRanks in the top 80th percentile of NHBE, A549, and BALF sample groups were each used as input for gProfiler:GOSt functional profiling ^22^. The input genes and resulting pathways were separated and used in downstream comparisons.

To explore co-regulated and influential components in the infection, genes with significant DE were correlated with SARS-CoV-2 transcripts for each group. Genes with an *R*^2^ relationship > 0.95 with a SARS-CoV-2 gene were used as input for gProfiler: GOSt functional profiling. The input genes and resulting pathways were separated and used in downstream analysis. To visualize the highest correlated elements, the ‘network’ package in R was used to plot an edge between a SARS-CoV-2 transcript and a gene if *R*^2^ > 0.98 ^45^. The size of the node in the network illustrates the degree of connection each node has relative to the other nodes in the network. It should be noted that, by virtue of the methodology, SARS-CoV-2 transcripts will have a higher degree of connectivity.

### Sankey Diagrams

To visualize how the four views affirm or disagree with one another, we created Sankey diagrams. For Figure 4B, the four views were connected to genes found in at least two of the four views. Genes were colored based on the views that they were found in: white, light blue, and dark blue indicate that DE and 1, 2, or 3 other views, respectively, and red for a gene that is not found by DE, but was by a CE view. These genes are then connected to themes, which consists of a list of gProfiler pathways that they were found to be enriched in (Suppl. Table 1). For Suppl. Figure 10, the four methods are connected to genes found by any of the four views and these genes.

### Network Construction

Finally, to visualize the gene and pathway agreement between the four views, the recorded genes and gProfiler pathways from SARS-CoV-2 infected groups were used to create gene to pathway networks for each view and for results that appear through multiple views. To be in these networks, an inclusion criteria was enforced where each gene and pathway displayed is found in at least two of the three SARS-CoV-2 infected sample types, namely BALF patient samples, NHBE infected cells, and A549 infected cells. An edge was created between genes and ontologies if they were implicated together by the gProfiler:GOSt result.

## Data Availability

All data used in this paper is available online ^11,12^.

