## Supplementary Figures and Legends for "Dual RNA-Seq analysis of SARS-CoV-2 correlates specific human transcriptional response pathways directly to viral expression"

**SUPPLEMENT**

### SUPPLEMENTAL FIGURES

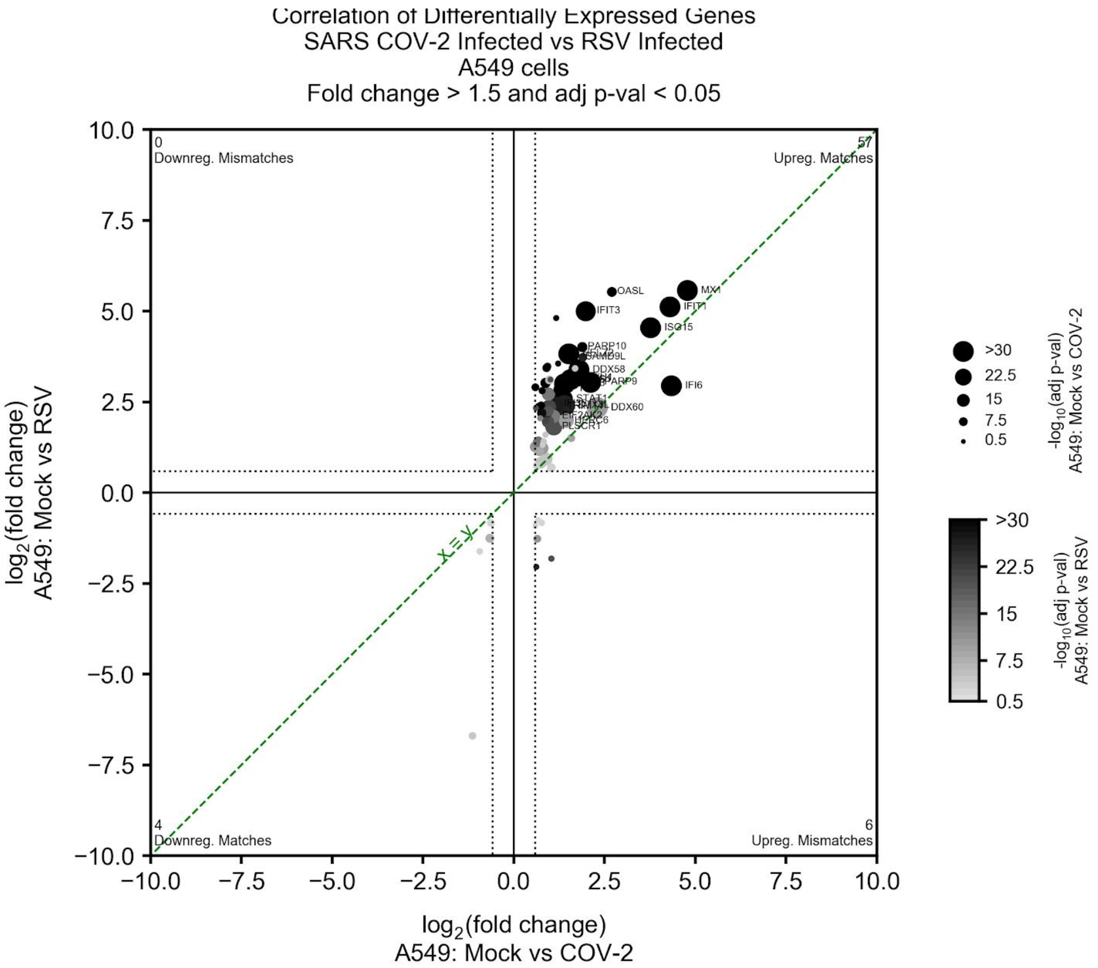


**Supplemental Figure 1.** Comparison differential expression programs in SARS-CoV-2 (COV-2) infected versus respiratory syncytial virus (RSV) infected cells. Differential expression (DE) of infected versus control in A549 cell lines produced a set of genes (circles, symbols displayed for a selected few) passing a log2 fold change (1.5) and significance cutoff (p-val<0.05, dashed vertical and horizontal lines) in both RSV (y-axis) and SARS-CoV-2 (x-axis) with high concordance (line of perfect correspondence shown; green line) as indicated by the number of genes with matching expression in the upper right quadrant of the plot (57 matches) or lower left (4 matches) compared to off-diagonal quadrants (0 and 6 genes). Significance levels of individual genes are depicted with size (COV-2) and intensity (RSV) from the two viruses.

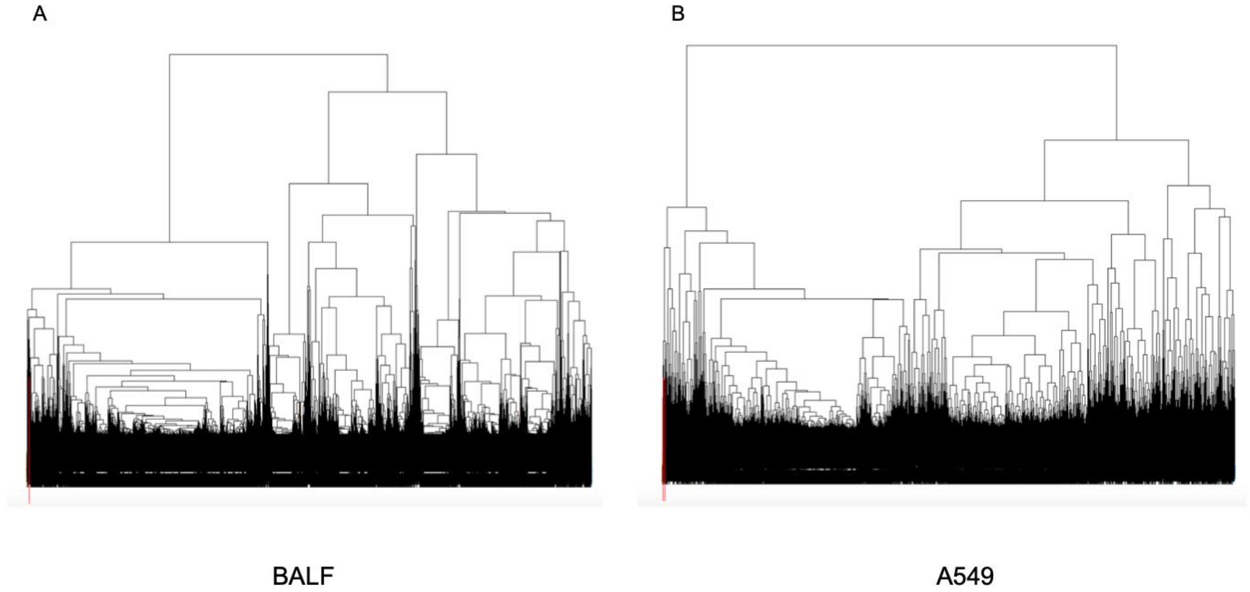


**Supplemental Figure 2.** Normalized gene counts are clustered by average linkage distance Pearson correlation and visualized by dendrograms. For both BALF tissue (A) and A549 cell lines (B), SARS-CoV-2 transcripts were located within a single clade (boxed in red to the left of the dendrogram) after 200 clades were created across the entirety of the hierarchical structure.

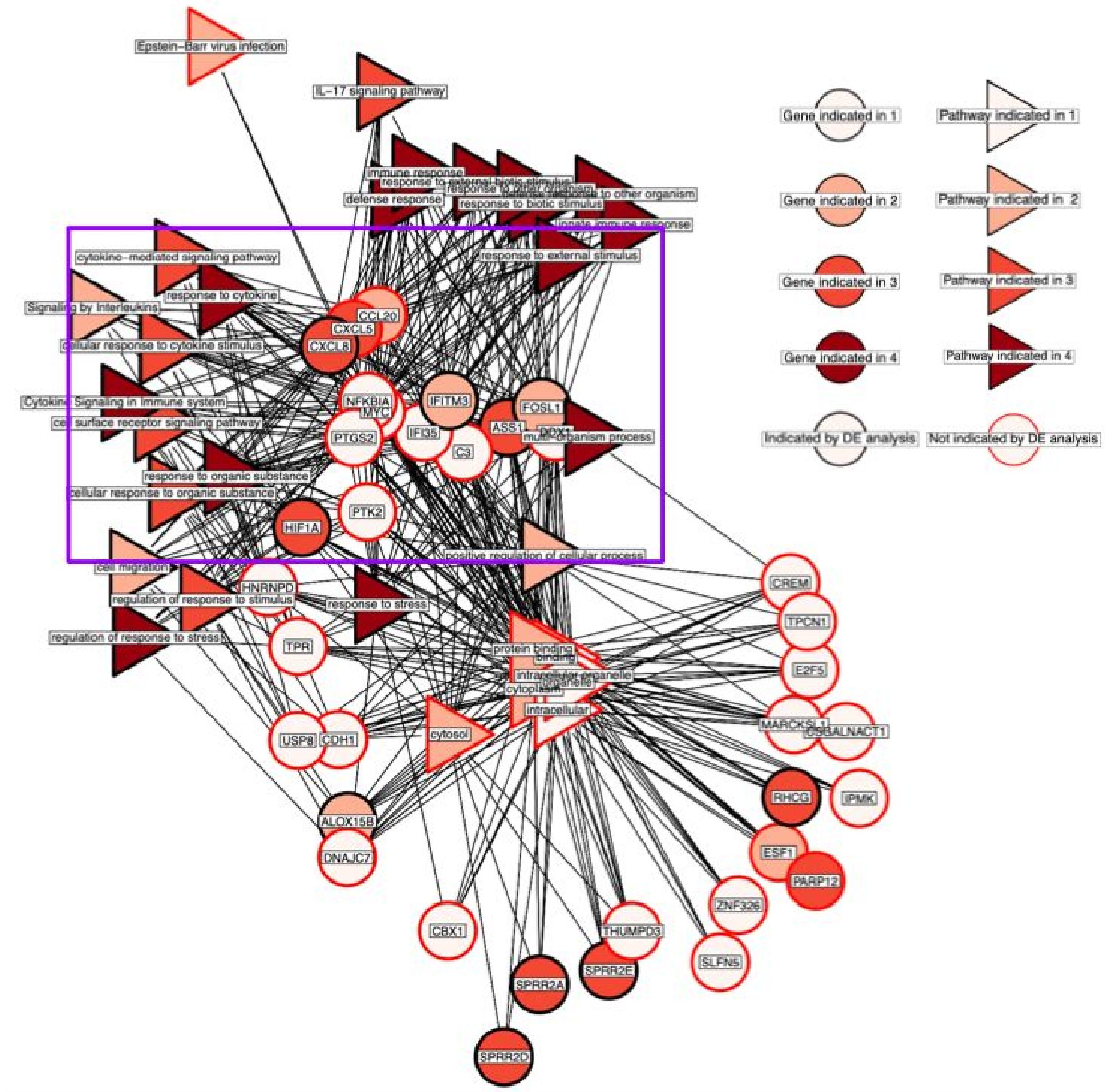


**Supplemental Figure 3.** The CE-Dendro gene and gProfiler pathways are visualized as a network where an edge is drawn between genes and the pathways for which they are members. A purple box is shown to highlight a particular group of genes with functional significance that were largely results not found by the DE view (CXCL5, CCL20, NFKBIA, MYC, PTGS2, IFI35, C3, DDX1, PTK2). These genes are important for cytokine and chemokine activity, antiviral response, and the innate immune system.

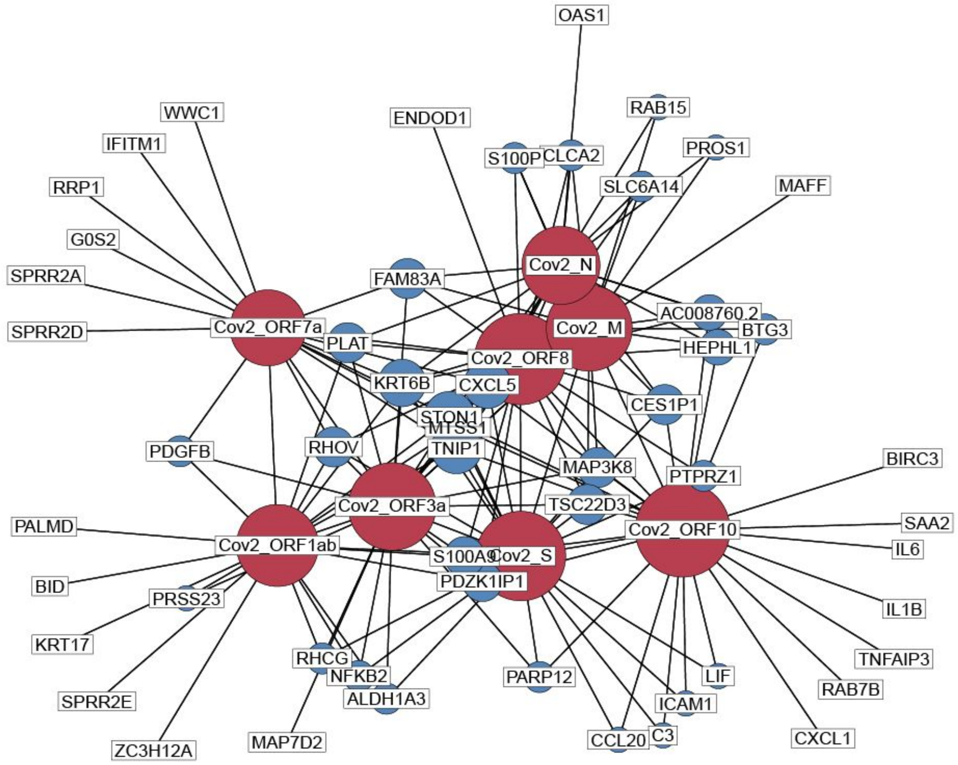


**Supplemental Figure 4.** Gene results from CE-Net in NHBE cells that were correlated with SARS-CoV-2 transcripts at |R| > 0.98 are visualized as a network where edges are drawn between human genes and the SARS-CoV-2 genes which they are correlated with. SARS-CoV-2 genes are shown in red and human genes in blue. Increasing node size indicates that the gene is connected to more nodes within the network. The S, ORF10, and ORF8 genes were found to be connected with the most human genes at this |R| threshold, suggesting that they may be functionally important for SARS-CoV-2 in its influence on the human transcriptome.

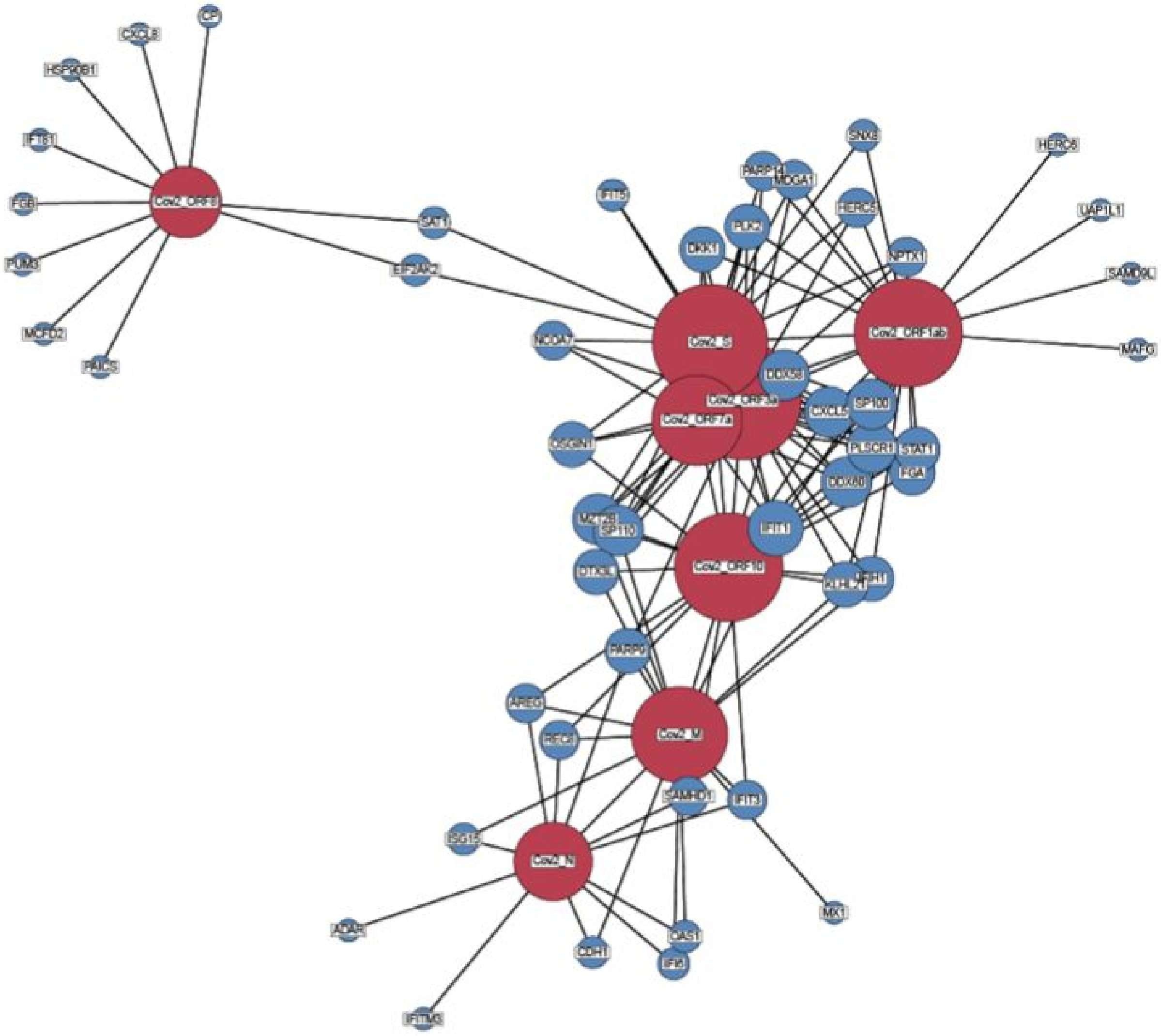


**Supplemental Figure 5.** Gene results from CE-Net in A549 cells that were correlated with SARS-CoV-2 transcripts at R > 0.98 are visualized as a network where edges are drawn between human genes and the SARS-CoV-2 genes which they are correlated with. SARS-CoV-2 genes are shown in red and human genes in blue. Increasing node size indicates that the gene is connected to more nodes within the network. The S, ORF10, ORF8, and ORF1ab genes were found to be connected with the most human genes at this |R| threshold, suggesting that they may be functionally important for SARS-CoV-2 in its influence on the human transcriptome.

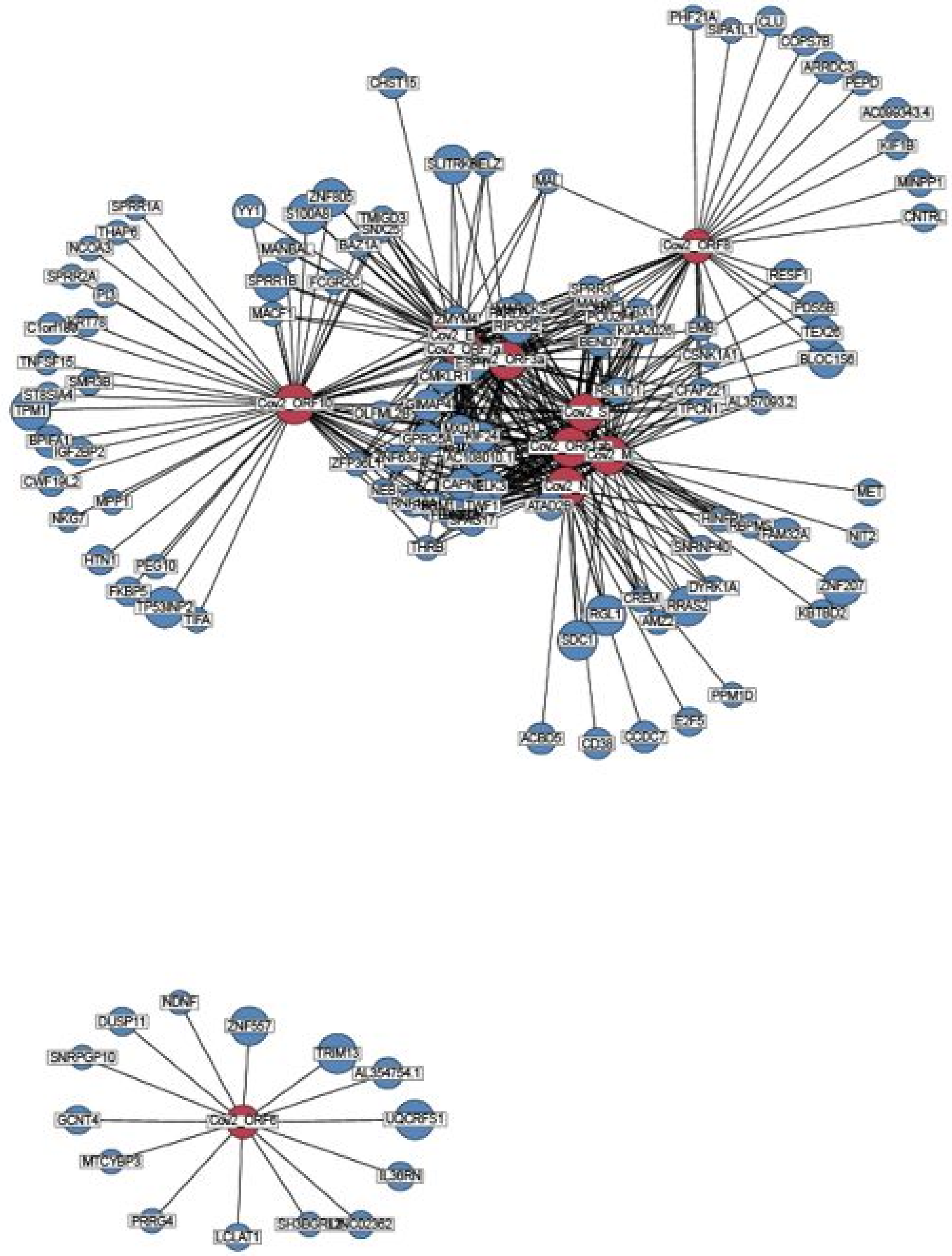


**Supplemental Figure 6.** Gene results from CE-Net in BALF tissue that were correlated with SARS-CoV-2 transcripts at R > 0.98 are visualized as a network where edges are drawn between human genes and the SARS-CoV-2 genes which they are correlated with. SARS-CoV-2 genes are shown in red and human genes in blue. Increasing node size indicates that the gene is connected to more nodes within the network. Most of the SARS-CoV-2 genes depicted in this network were similarly connected to many human genes. In contrast, ORF6 was not co-correlated to human genes with any other SARS-CoV-2 gene.

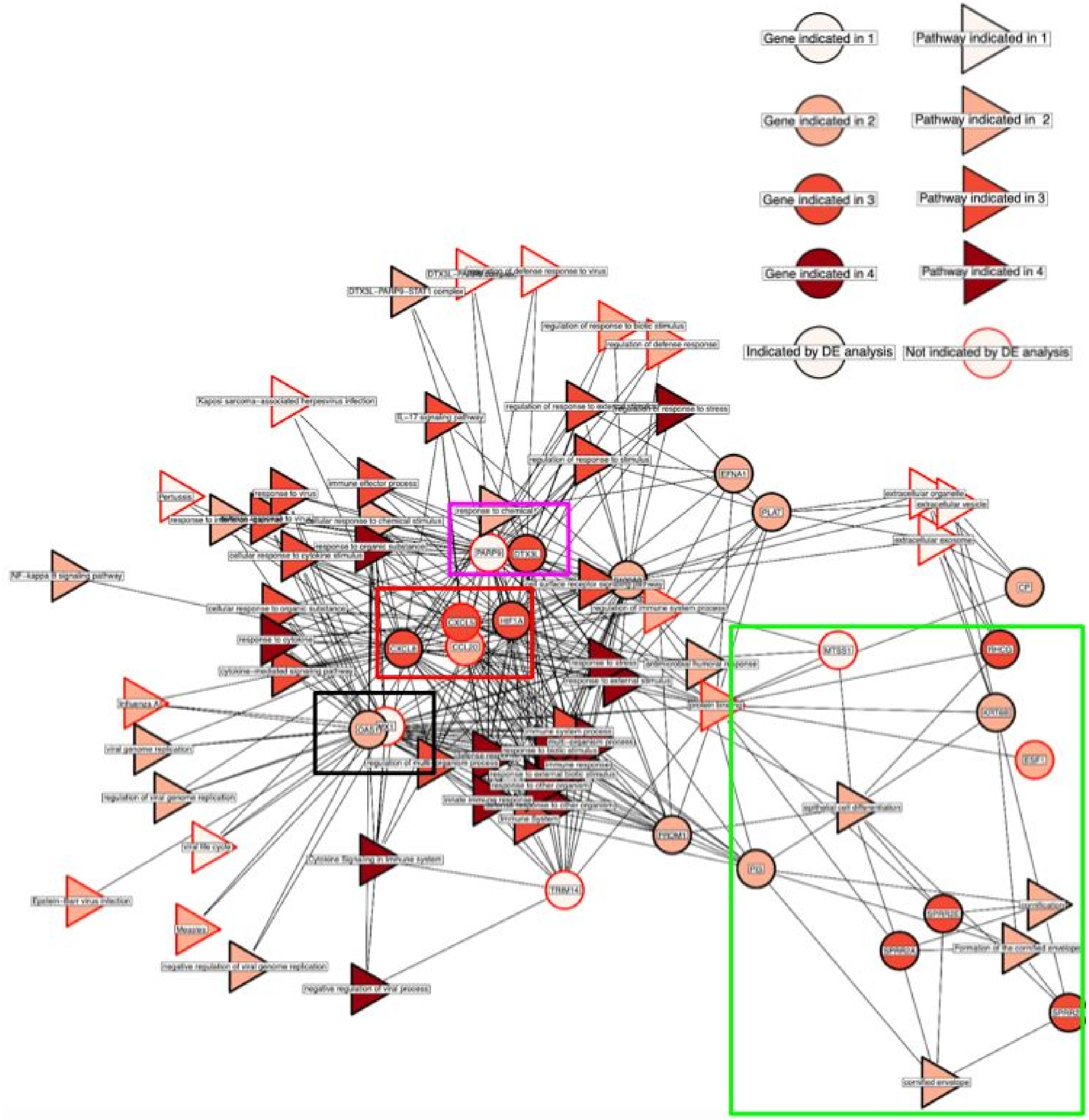


**Supplemental Figure 7.** The CE-Net gene and gProfiler pathways are visualized as a network where an edge is drawn between genes and the pathways for which they are members. The black box shows genes (OAS1 and MX1) with antiviral properties. The red box shows genes indicative of chemokine and cytokine activity (CXCL5, CXCL8, CCL20, HIF1A). The pink box highlights the PARP9-DTX3L gene group. Finally, the green box shows a group of genes involved in cornification and cell death processes (MTSS1, RHCG, KRT6B, ESF1, PI3, SPRR2A, SPRR2D, SPRR2E).

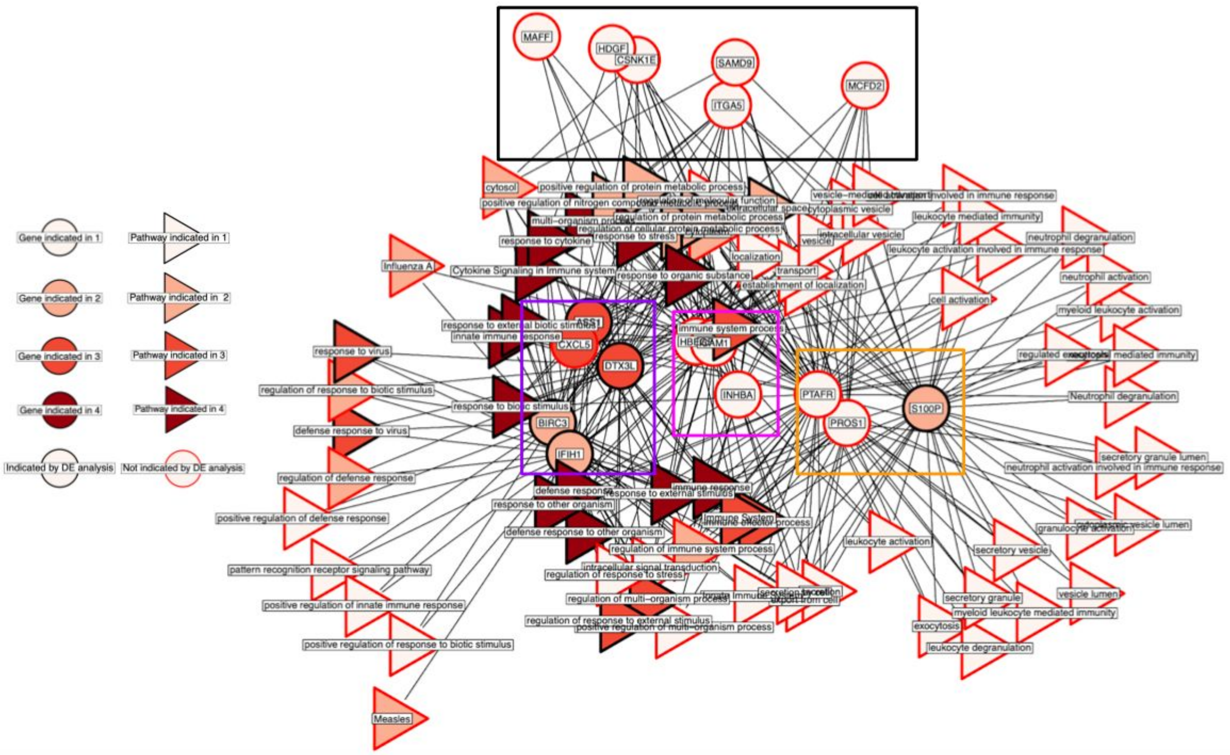


**Supplemental Figure 8.** The CE-PageRank gene and gProfiler pathways are visualized as a network where an edge is drawn between genes and the pathways for which they are members. The purple box highlights a gene group (ASS1, CXCL5, DTX3L, BIRC3, IFIH1) indicative of lipopolysaccharide response and cytokine, chemokine, and immune activity. The pink box shows genes (ICAM1, HBEGF, INHBA) involved in various processes including viral entry and survival. The yellow box shows genes involved in calcium binding and secretory activity (S100P, PROS1, PTAFR). Lastly, the black box highlights gene results that were not found by DE (MAFF, HDGF, CSNK1E, ITGA5, SAMD9, MCFD2).

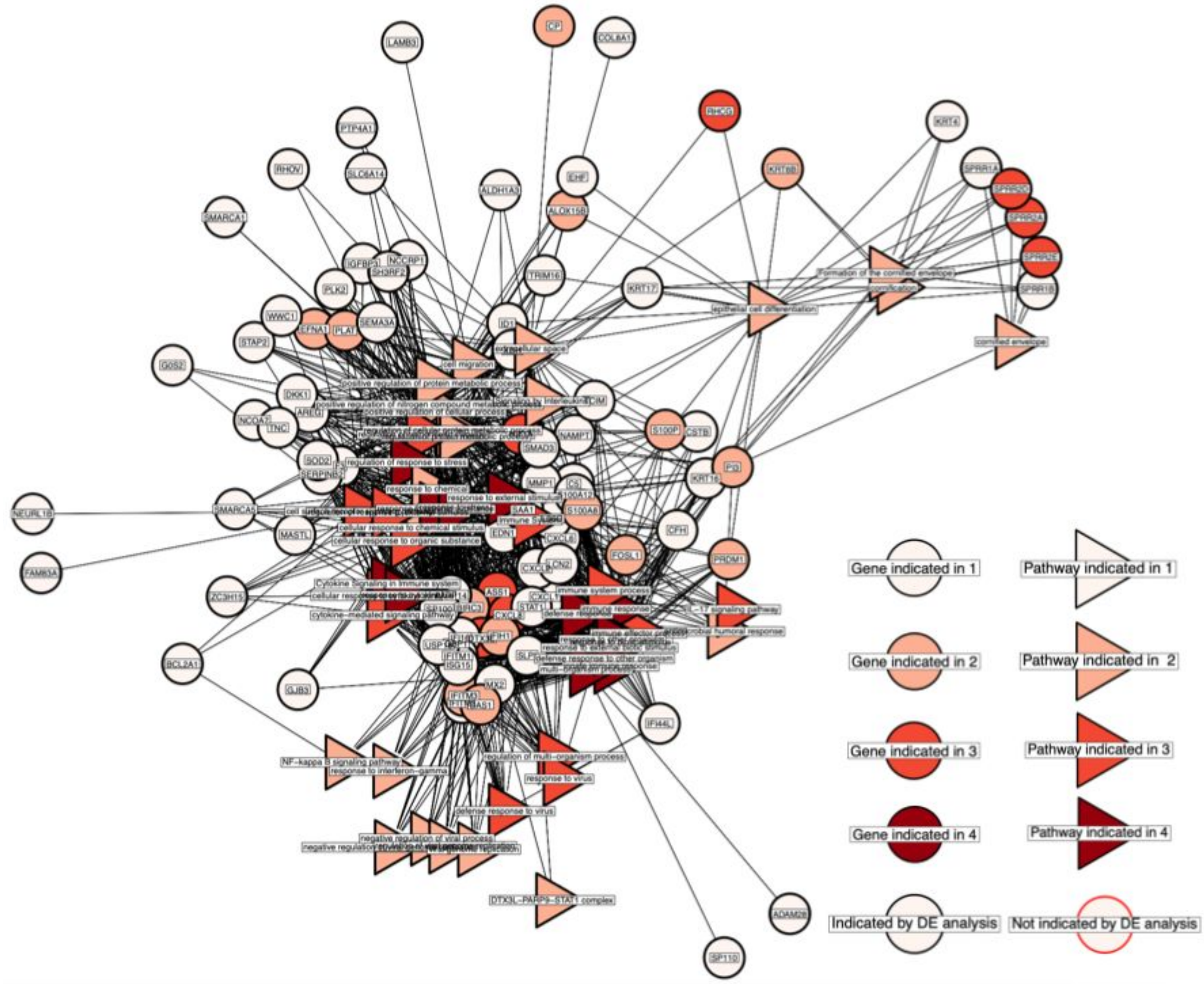


**Supplemental Figure 9.** The DE gene and gProfiler pathways are visualized as a network where an edge is drawn between genes and the pathways for which they are members. White nodes (indicating genes or pathways that were only found by DE) were a dominant feature (72 genes) among the gene results for DE, suggesting that DE highlights the conditional response of human genes to SARS-CoV-2 infection.

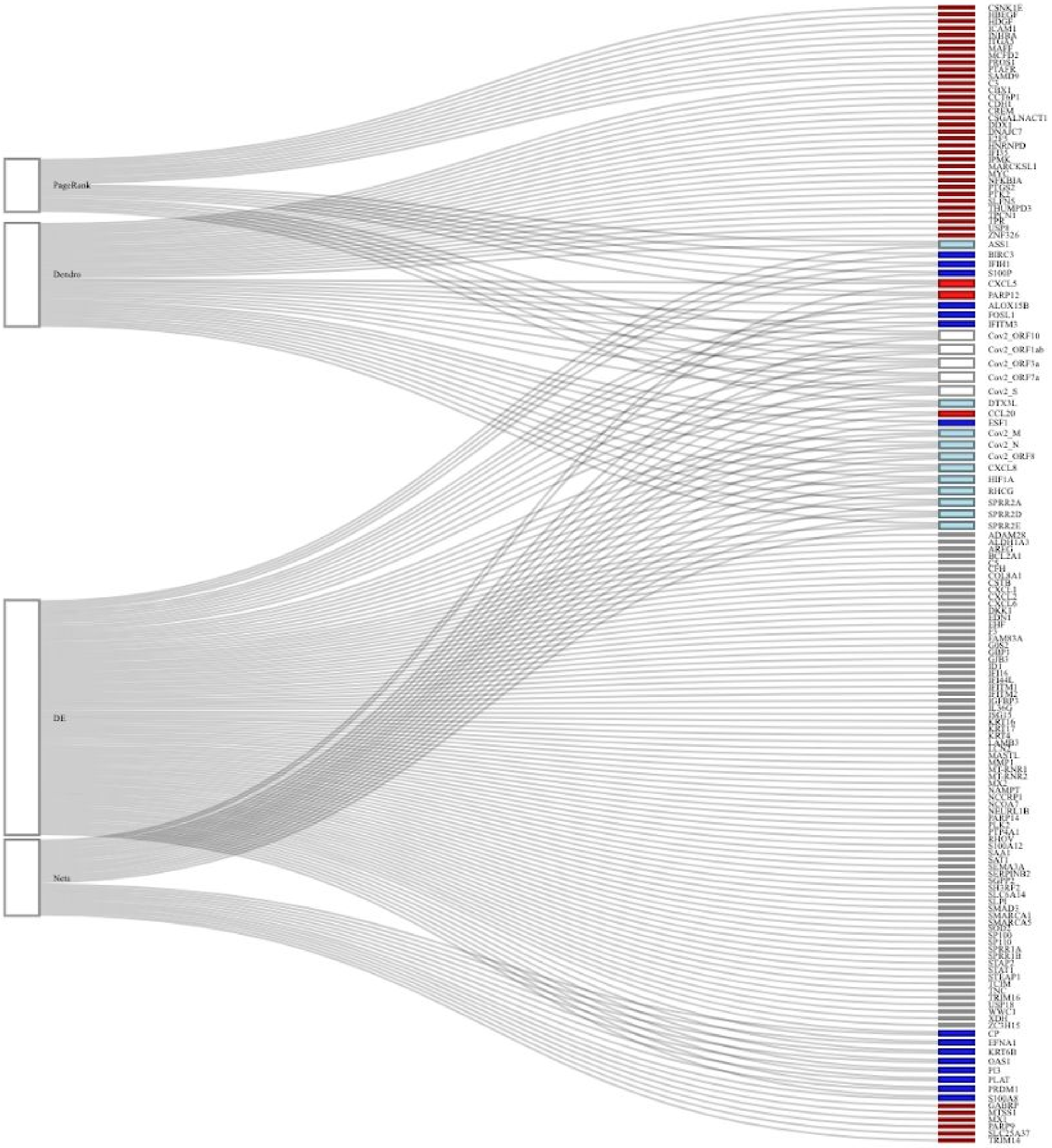


**Supplemental Figure 10.** The sankey diagram shows genes that were found by any of the four views and the pathway themes that they participate in. Genes are colored based on the views that they are found in, white, light blue, and dark blue indicate that DEA and 1, 2, or 3 other views, respectively, and red for a gene that is not found by DEA, but was by a CEA view.
